# South Asian Patient Population Genetics Reveal Strong Founder Effects and High Rates of Homozygosity – New Resources for Precision Medicine

**DOI:** 10.1101/2020.10.02.323238

**Authors:** Jeffrey D. Wall, J. Fah Sathirapongsasuti, Ravi Gupta, Anamitra Barik, Rajesh Kumar Rai, Asif Rasheed, Venkatesan Radha, Saurabh Belsare, Ramesh Menon, Sameer Phalke, Anuradha Mittal, John Fang, Deepak Tanneeru, Jacqueline Robinson, Ruchi Chaudhary, Christian Fuchsberger, Lukas Forer, Sebastian Schoenherr, Qixin Bei, Tushar Bhangale, Jennifer Tom, Santosh Gopi Krishna Gadde, B. V. Priya, Naveen Kumar Naik, Minxian Wang, Pui-Yan Kwok, Amit V. Khera, B. R. Lakshmi, Adam Butterworth, John Danesh, Sekar Seshagiri, Sekar Kathiresan, Arkasubhra Ghosh, V. Mohan, Abhijit Chowdhury, Danish Saleheen, Eric Stawiski, Andrew S. Peterson

**Author notes:** equal contributions (Jeffrey D. Wall, J. Fah Sathirapongsasuti, Ravi Gupta, Anamitra Barik, Rajesh Kumar Rai, Asif Rasheed, Venkatesan Radha). equal contributions (V. Mohan, Abhijit Chowdhury, Danish Saleheen, Eric Stawiski, Andrew S. Peterson).

## Abstract

Population-scale genetic studies can identify drug targets and allow disease risk to be predicted with resulting benefit for management of individual health risks and system-wide allocation of health care delivery resources. Although population-scale projects are underway in many parts of the world, genetic variation between population groups means that additional projects are warranted. South Asia has a population whose genetics is the least characterized of any of the world’s major populations. Here we describe GenomeAsia studies that characterize population structure in South Asia and that create tools for economical and accurate genotyping at population-scale. Prior work on population structure characterized isolated population groups, the relevance of which to large-scale studies of disease genetics is unclear. For our studies we used whole genome sequence information from 4,807 individuals recruited in the health care delivery systems of Pakistan, India and Bangladesh to ensure relevance to population-scale studies of disease genetics. We combined this with WGS data from 927 individuals from isolated South Asian population groups, and developed a custom SNP array (called SARGAM) that is optimized for future human genetic studies in South Asia. We find evidence for high rates of reproductive isolation, endogamy and consanguinity that vary across the subcontinent and that lead to levels of homozygosity that approach 100 times that seen in outbred populations. We describe founder effects that increase the power to associate functional variants with disease processes and that make South Asia a uniquely powerful place for population-scale genetic studies.

---

Founder effects and population bottlenecks reduce the number of individuals from the past that contribute to present day genetic diversity. The shifts in allele frequencies that result have contributed to many important discoveries in studies of Icelandic, Ashkenazi, Finnish, Amish and other founder or bottlenecked populations^1-4^. The historical events that have produced genetic drift in these populations are recognizable and the genetic consequences can be effectively modeled. Studies of population structure in South Asia have described patterns of genetic drift as founder effects^5^ but there is little evidence that reductions in population size have been a significant factor in producing present day population structure. Endogamy (i.e., marriages that are restricted to a particular group or caste) is however well recognized in South Asia^6^ and can, through effective reproductive isolation, produce founder effects by reducing *effective* population size. At one extreme is consanguineous partnering but there are many possible reproductive patterns that can be described as endogamy, and it is not possible to usefully predict all of the possible effects on genetic variation. Empirical description of population structure as it presents to the health care delivery system has been lacking but is critical in order to efficiently design and carry out human genetic studies at population scale. At the same time, tools for accurate and economical genotyping, also necessary for population scale genotyping, have been lacking for South Asians. Used together, a clear description of population structure and economical genotyping tools can unlock the tremendous potential of human genetics in South Asia for discoveries that illuminate disease processes and allow prediction of disease risk in South Asians.

## The GAsP2 data set

We divided samples from the health care delivery systems of South Asia (informally called ‘medical cohorts’) into three regional groups: Pakistani (PAK), South Indian (SOI) and Bengali (BNG). We combined these samples with previously published genomes^7-12^ and additional samples we sequenced from isolated South Asian population groups to create the GenomeAsia Phase 2 (GAsP2) data set. We then used a standard pipeline for read mapping and variant calling, starting from the raw sequence data of all of the samples. After standard quality control filters and removal of one individual from each first-degree relative pair, we obtained a set of 6,443 high-coverage genomes (average 25x) for downstream analyses (Supplementary Table 1). Of these, 5,734 genomes were of South Asian ancestry, with medical cohort sizes of 1,810, 1,363 and 1,634 respectively for the Pakistani, South Indian and Bengali groups. Basic information on SNPs and allele frequencies are available from the GenomeAsia consortium website (https://www.genomeasia100k.org), while a computationally phased version of the data can be used as a reference panel for imputation using the Michigan Imputation Server (https://imputationserver.sph.umich.edu/index.html). The estimated non-reference discordance rate for duplicate samples found in both the GAsP2 and 1000 Genomes Project data sets is 1.61 * 10^−4^ (Supplementary Table 2).

## Population structure

We used standard approaches such as F_ST_, PCA^13^, Admixture^14^ and UMAP^15^ (Uniform Manifold Approximation and Projection) for qualitatively assessing population structure in our study (Fig. 1, Supplementary Figures 1-4) In the UMAP plot of South Asian samples, we find a clear distinction between Bengali, Pakistani and South Indian groups that roughly mirrors geography (Fig. 1a) When we further zoom in on each of these three regions, we find that UMAP can separate individuals into smaller caste and culture-based subgroups (Fig. 1b-d) In particular, since our samples include both medical cohorts (sampled from particular regions without regard to caste) and focused sampling of particular caste and language groups, we can often reliably assign caste (Fig. 1d) or subgroup (Fig. 1b) labels to individuals based solely on their genetic makeup.

**Figure 1.**
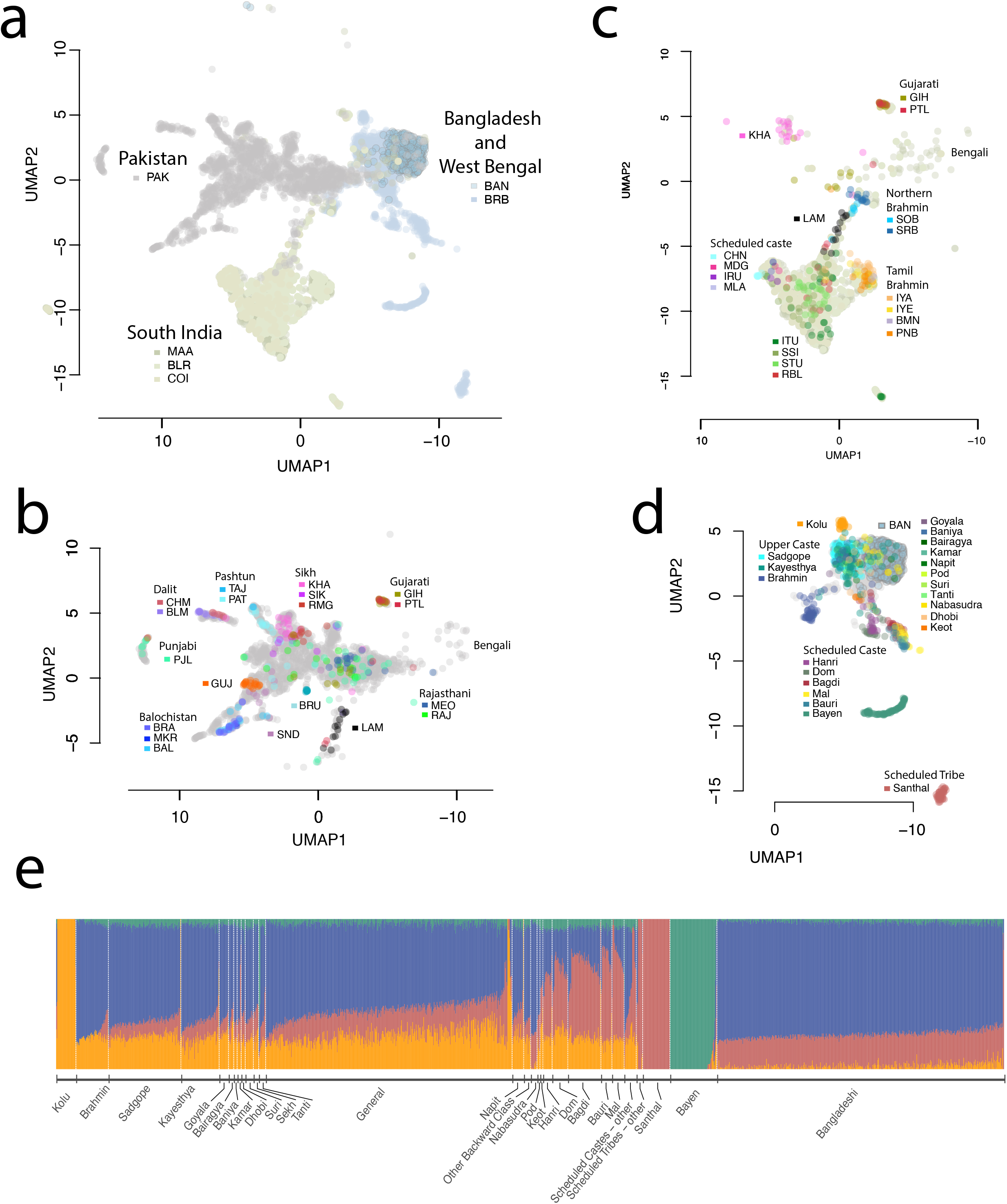
Fine-scale population structure in the Health-care delivery system. UMAP was run on all samples using the first 15 principal components. **a**, In the South Asian subset, samples cluster into three major groups by sample origins: Pakistan; South India; and West Bengal and Bangladesh. The X-axis (UMAP1) was flipped along the vertical axis so that the parallel between the graphical position of the three populations and the map of South Asia was apparent. **b, c**, Samples with detailed locations or self-reported group memberships are shown to segregate within Pakistan and South India clusters. Among the samples from Pakistan and South India, some segregate with recent immigrants (e.g. Bengalis and Gujaratis) and historical immigrants (e.g. Lambadas), reflecting the metropolitan nature of the recruitment centers. **d**, Samples from Birbhum District, West Bengal, have detailed self-reported group membership information. Upper castes, scheduled castes, and scheduled tribes clearly segregate, reflecting the historical reproductive isolation between these groups. Bayen and Santhal are two notable population isolates. **e**, ADMIXTURE analysis of samples from the Birbhum District shows four major components. Labels are self-reported group identity with “general” denoting a lack of specified identity. PAK - Pakistan; BLR - Bangalore; MAA - Chennai; COI - Coimbatore; BAN - Bangladesh; BRB - Birbhum District, West Bengal; LAM - Lambada. For other 3-letter codes, see Supplementary Table XX.

In Birbhum, West Bengal, we have self-reported population group identity (e.g. tribe, caste and/or sub-caste) for over half of the individuals in our study. Figure 1c shows that some clearly reported population group identities are genetically distinct, such as the Santhal, Bayen and Brahmin, while others are not, like the Sadgope and Kayastha. An admixture plot of Birbhum and Bangladeshi samples with K = 4 provides a similar picture (Fig. 1e), with Santhal, Bayen and Kolu appearing to be well-defined genetic groups, while most individuals from the other groups are estimated to be admixed. The clustering of the Santhal, Bayen and Kolu reflects increased genetic drift due to some combination of isolation, endogamy and consanguinity. It is well established that the increased drift in founder populations in general and South Asian groups in particular enable unique opportunities for genetic research^5,16^. Among the other groups, we note that there is a clear distinction in Admixture estimates between Bangladeshis and general caste individuals from Birbhum (who include a substantial number of Bengali Muslims). Since Partition (between India and what was then East Pakistan) happened too recently to cause systematic subsequent genetic differences, our results suggest that the residents of Bangladesh are a non-random subset of the Muslims that were living in Bengal in the mid-20^th^ century.

## Endogamy and consanguinity

Endogamy and consanguinity lead to an excess of homozygous genotypes over the expectations from random-mating. We calculated the ratio of the observed number of rare homozygous genotypes over the expected number, binned by minor allele frequency (MAF), for South Indian (SOI), Pakistani (PAK) and Bengali (BNG) samples (Fig. 2a). For comparison with the patterns of genetic variation in an ostensibly outbred population, we also included the same results for 1,442 unrelated Taiwanese (TWN) genomes. For all four groups, we observe an increasing excess of rare homozygotes with smaller MAF. This pattern is strongest in the South Indian and Pakistani groups, moderate in the Bengali group, and weak in the Taiwanese, and reflects the strength of non-random mating within each group.

**Figure 2.**
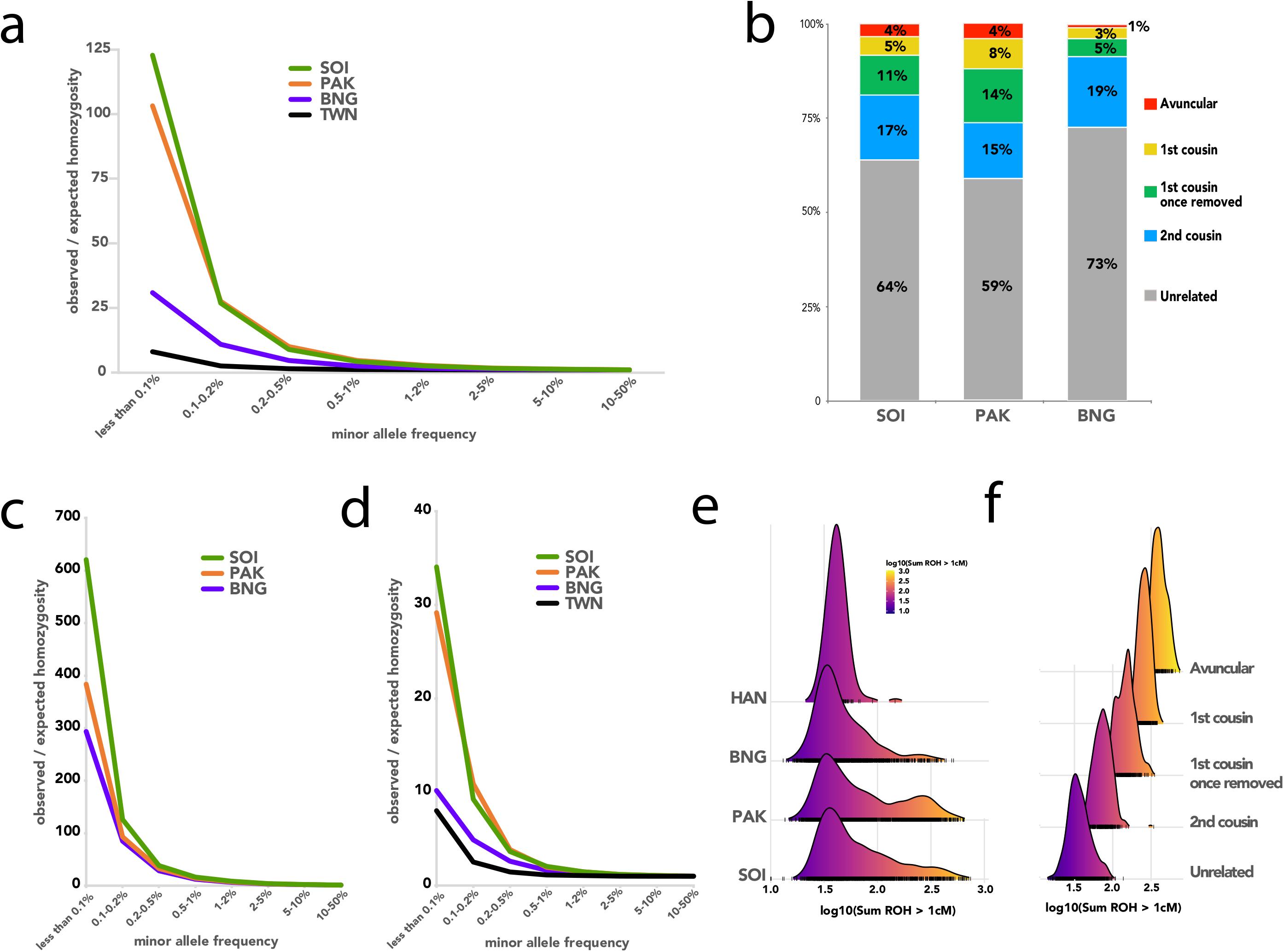
Homozygosity and inbreeding in South Asian cohorts, **a**. Observed / expected proportions of rare homozygotes, stratified by minor allele frequency and population. The expected values assume random mating, **b**. Stacked bar chart showing the estimated degree of inbreeding for individuals in the South Asian medical cohorts, **c**. Same as in panel a but calculated for “inbred” individuals (whose parents are estimated to be 3rd degree relatives or more closely related) only. **d**. Same as in panel a but calculated for “outbred” individuals (whose parents are estimated to be 6th degree relatives or more distantly related) only. **e**. Ridgeplots showing the distribution across individuals of the total (genetic) length of the genome contained in ROHs that are at least 1 cM in length, **f**. Ridgeplots showing the stratification of Fig. 2e’s PAK plot into groups with different estimated degrees of inbreeding.

Founder events or population bottlenecks that occurred in the distant past can dramatically increase the rate of homozygosity, but with tract lengths decreasing over increasing numbers of intervening generations. Endogamy and consanguinity both produce excess homozygosity, but in the latter the excess homozygosity primarily occurs in long runs of homozygosity (ROH). To determine the relative effects of endogamy and consanguinity on patterns of homozygosity within our South Asian cohorts, we developed a novel method for estimating the degree of parental relatedness of an individual based on the observed numbers and lengths of long (e.g., >10 cM) ROHs. The method categorized an individual’s parents as 2^nd^ degree relatives (e.g., uncle and niece), 3^rd^ degree relatives (e.g., 1^st^ cousins), 4^th^ degree relatives (e.g., 1^st^ cousins once removed), 5^th^ degree relatives (e.g., 2^nd^ cousins) or unrelated (i.e., less related than 2^nd^ cousins). Both the proportion of individuals identified as outbred and the distribution of consanguineous individuals across the remaining four categories show substantial regional variation (Fig. 2b). In particular, there appears to be less consanguinity and fewer closely related parental pairings on average among the individuals in BNG (West Bengal and Bangladesh), compared with the medical cohorts from SOI (South India) and PAK (Pakistan). This presumably reflects systematic differences in marriage practices across the different regions. Interestingly, self-reported consanguinity in Birbhum samples is only modestly correlated with genetic estimates of consanguinity (Supplementary Figure 5).

To assess whether consanguinity by itself can explain the observed excess of rare homozygotes, we stratified each regional group into ‘inbred’ and ‘outbred’ subgroups (where the former referred to individuals whose parents were estimated to be 2^nd^ or 3^rd^ degree relatives). We then tabulated the increase in rare homozygotes for each subgroup (Fig. 2c and d). We find that the inbred subgroups (Fig. 2c) have as much as 600 fold higher levels of rare homozygotes above expectation, 4-10 fold above the population group as a whole and 13-30 times the level seen in their outbred sub-group. Interestingly, even in the outbred South Asian sub-groups (Fig. 2d) levels of homozygosity can be as much as 3.8 fold higher than in TWN (Han Chinese from Taiwan) individuals, presumably due to endogamy and the resulting enrichment in distant parental relationships for the individuals in our data set which our methods were not able to identify.

Excess homozygosity caused by close parental relatedness is structured in long runs of homozygosity (ROH). We tabulated the total length of each individual’s genome contained in ROH longer than 1 cM, and plotted the distribution of this length across several South Asian groups, along with HAN (Han Chinese from China) as an outbred population for comparison (Fig. 2d). All of the South Asian groups have a tail of individuals with increased proportion of ROH that corresponds with the individuals that have parents that are closely related (Fig. 2e). Consistent with this interpretation, subdividing the medical cohort samples based on parental relatedness deconvolutes the tail into separate distributions of ROH (Fig. 2f and Supplementary Figure 6).

## Loss of Function variants

To assess the potential functional effects of the high levels of endogamy and consanguinity found within South Asia, we identified putative Loss of Function (pLoF) variants in our data set (see Methods). For comparison, we also used an analogous analysis pipeline to identify pLoF variants in non-Finnish European (NFE) individuals from the genome aggregation database^17^ (gnomAD). There are more genes with pLoF variants with frequency > 0.1% found in South Asia (SAS) than in NFE, and most of these genes are in the set of pLOF containing genes unique to SAS (Fig. 3a). To visualize the frequencies and geographic distributions of these pLoF variants, we constructed heat maps representing genes containing pLoF variants, with warmer colors indicating higher minor allele frequencies of pLoF variants. Individual clusters are shown for each set of pLOF containing genes that are unique to or shared between individual population groups (Fig. 3b). The excess of pLOF containing genes that is seen in SAS is distributed fairly evenly amongst PAK, BNG and SOI, consistent with the idea that these are distinct population groups, each of which had founder effects that pushed LOF and other functionally relevant alleles to higher frequencies than is seen in an outbred population. We also show the same heat maps for homozygous pLoF variants (i.e., human ‘knockouts’) in Supplementary Figure 7.

**Figure 3.**
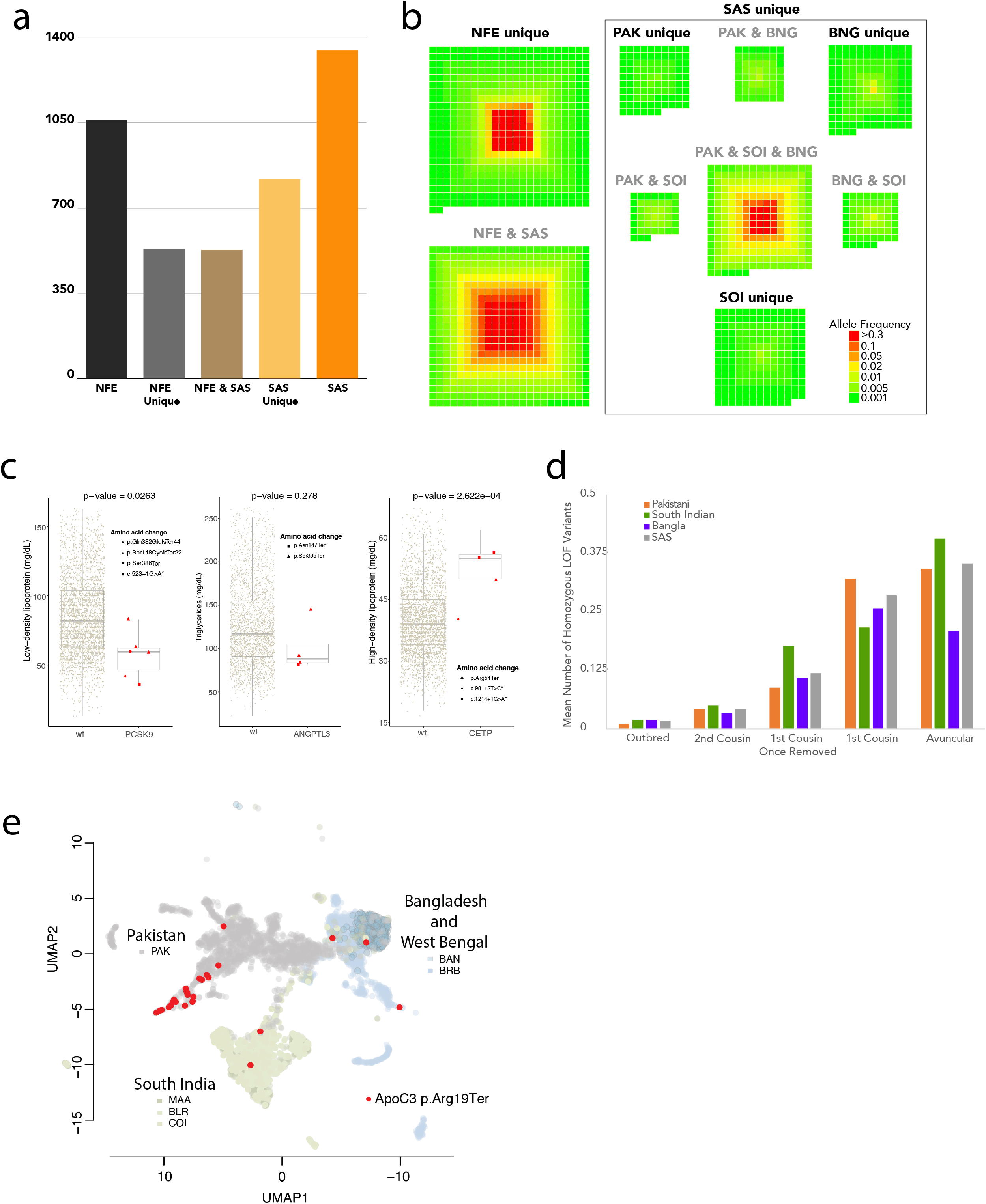
Loss of function mutations. **a**, Number of high confidence loss of function genes found at a minimum of 0.1% AF in their relative population: total found in non-Finnish European (NFE), found in NFE and not in SAS (NFE Unique), found in both NFE and SAS (NFE & SAS), found in SAS and not NFE (SAS Unique) and total found in SAS. **b**, Loss of function gene space by population. Each square represents a distinct gene and is colored by its maximum AF within the relative group. Genes are separated by groups in which they are found (from top to bottom and then left to right): NFE unique, NFE & SAS, PAK unique, PAK & SOI, PAK & BNG, all of SAS (PAK & SOI & BNG), SOI unique, BNG unique, and BNG & SAS. **c**, Effects of pLoF variants on blood lipid markers: PCSK9 pLoFs associated with decreased LDL, ANGPTL3 pLoFs associated with decreased triglyceride, and CETP pLoF associated with increased HDL. Only samples from South India (Bangalore and Chennai) were included. **d**, Mean number of homozygous pLoF variants per individual, stratified by population and estimated degree of inbreeding. **e**, APOC3 p.Arg19Ter alleles are found at a high frequency among Balochi and Sindhi individuals in Southern Pakistan. Three of the identified Balochis and Sindhis were heterozygous carriers, and a larger number of carriers without self-reported identity were mapped to the same locus on the UMAP plot.

pLoF mutations are widely studied because they often have phenotypic effects that can easily be tied to the function of a specific gene or pathway. We looked at three genes where loss-of-function mutations are known to affect blood lipid levels^18-20^ and verified that individuals in our study that have pLoF variants in these genes have the expected effects on measured LDL, triglyceride and HDL levels (Fig. 3c).

A recognized benefit of studying South Asian populations is the greater probability of identifying individuals homozygous for pLoF alleles due to the excess homozygosity caused by endogamy and consanguinity. To evaluate this potential explicitly in our dataset, we tabulated the average numbers of rare (MAF < 0.01) homozygous pLoF mutations per individual (i.e., those pLoF mutations most likely to be deleterious), stratified by estimated degree of consanguinity (Fig. 3d). As expected, increased consanguinity is associated with an increased number of these rare, likely harmful mutations, similar to previous findings (cf. Figure 1c in ref. 21). As the degree of consanguinity increases, these mutations are more likely to be found in long ROH caused by recent inbreeding (Supplementary Figure 8).

The population structure within South Asia makes the region ideal for prospective studies of loss-of-function mutations. Even rare pLoF variants might have appreciable frequency in particular regions or caste groups, which would enable focused recruiting for follow-up functional studies. To evaluate this, we displayed the distribution of individuals containing characterized LOF alleles on the UMAP plots described previously (Fig. 3e and Supplementary Figure 9). Characterization of ApoC3 LOF homozygotes has elucidated the physiological basis by which ApoC3 acts to regulate serum triglycerides^21^. Within South Asia, ApoC3 LOF carriers are found predominantly in Pakistani subpopulations that cluster with individuals from Balochistan and Sindh in the South of Pakistan (Fig. 3e and 1b).

## The SARGAM Genotyping SNP array

To optimize the effectiveness of future genotype-phenotype studies in South Asia, we worked with Thermo Fisher to design a custom SNP array (South Asian Research Genotyping Array for Medicine, or SARGAM) that (i) prioritizes direct genotyping of known or putative protein altering variants present at a frequency of 0.1% or higher in SAS populations, and (ii) is optimized for imputation of variants down to an SAS minor allele frequency of 0.1%. To highlight the former feature, we tabulated how many pLoF or presumed damaging mutations can be directly genotyped by the most commonly used technology at present, Illumina’s GSA3 array and by the SARGAM array (Fig. 4a-b, Supplementary Figure 10). In Figure 4a and b, each protein coding gene is represented by a square in an array of 19,600 squares, with each square colored by the number of deleterious variants captured by each array at each gene. The SARGAM array directly genotypes presumed damaging mutations from the vast majority (74%, n=14,713) of non-read through protein coding genes in Ensembl (Fig. 4a), with a mean coverage of 3.5 mutations per gene (n=51,804). In contrast, the number of damaging mutations genotyped by the GSA3 array is much smaller (Fig. 4b), covering just 26% of genes (n=5,100) with a mean coverage of 1.9 mutations per gene (n=9,443). The SARGAM array therefore represents an inexpensive method for simultaneously conducting many specific genetic tests, while also allowing for standard human genetic applications (e.g., genome-wide association studies and/or polygenic risk score calculations).

**Figure 4.**
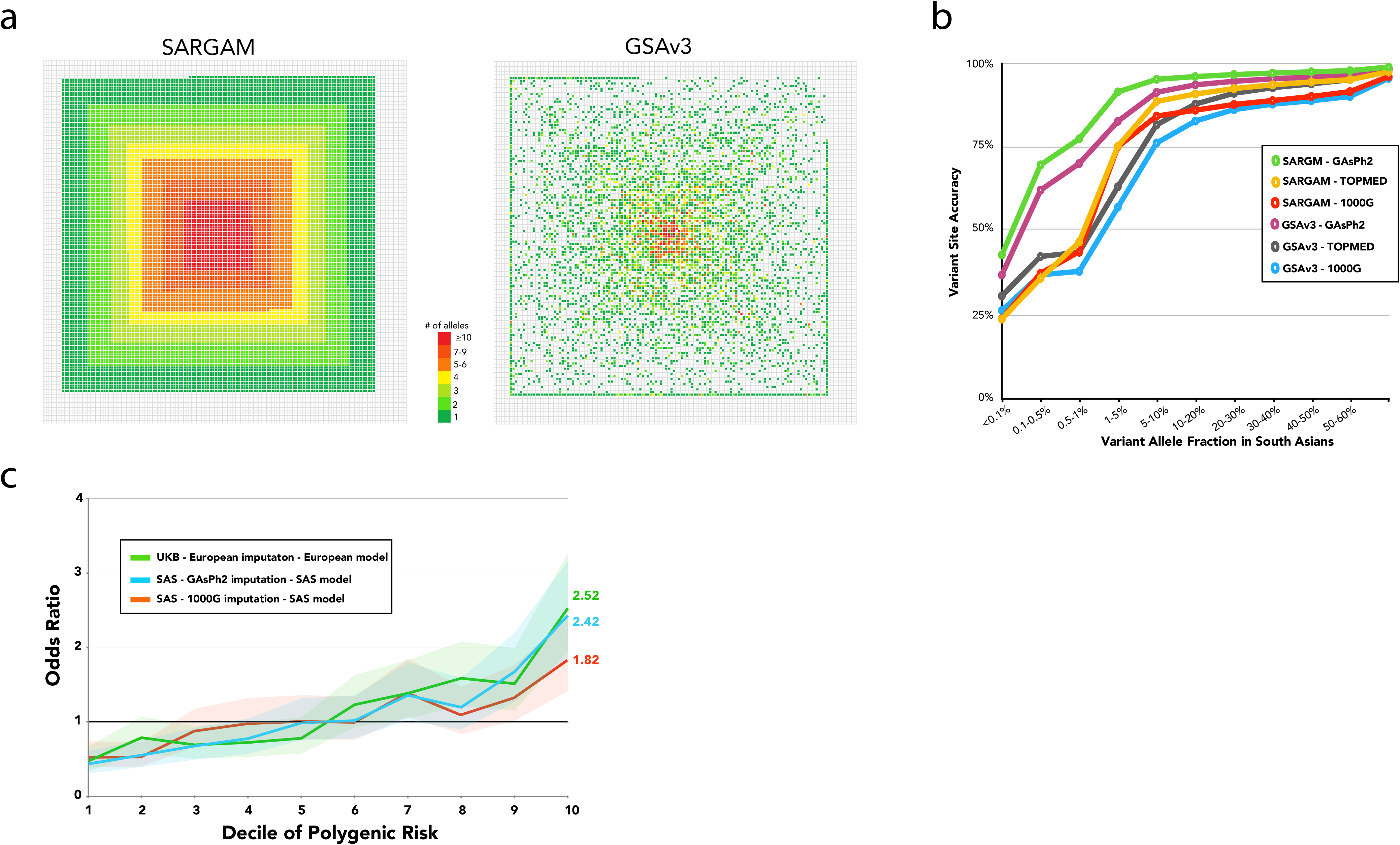
Improved Genotyping of South Asian Genomes. **a**. Gene space plot of all protein altering alleles that are directly genotyped using either the SARGAM or the Illumina GSA3 arrays. Protein coding genes of the human genome are depicted as an array of 19,600 squares. Genes with genotyped variants are shown as squares that are colored to indicate the number of gene-specific variants that are genotyped. **b**. Accuracy of non-reference allele imputation for combinations of genotyping array and imputation reference panel. Array genotypes were modeled by down-sampling from an independent data set of 30x WGS data. Missing genotypes were imputed using the indicated reference panels and the variant site accuracy of non-reference alleles was calculated and graphed as a function of allele frequency in South Asian genomes, **c**. Impact of imputation on Polygenic Risk Score (PRS) calculation. PRS were calculated using imputed genotypes from a CAD case-control cohort of 3,000 South Asian individuals genotyped using the Illumina GSA3 array and using a SAS PRS model (Wang et al 2020). The individuals were divided into 10 groups based on deciles of PRS and odds ratios were calculated from case-control status of the individuals in each group. For comparison, a case-control cohort of white Britons, matched for age and gender with the SAS cohort, was selected from the UKBiobank data set. PRS was calculated using a European model (Khera et al, 2018). 95% confidence intervals are shown for each PRS result as a shaded area.

Since the SARGAM array design utilized the observed patterns of linkage disequilibrium in thousands of South Asian genomes, we expected it to allow for more accurate imputation of untyped genotypes in South Asian samples. To evaluate this we compared imputation accuracy between the SARGAM and GSA3 arrays and found that both the SARGAM array and the GAsP2 reference panel contribute to higher imputation accuracy (Fig. 4c). These results reinforce why the quality of available genomic resources is a factor that limits feasibility and power of large-scale human genetic studies in non-European populations.

## Polygenic risk scores and the genetic architecture of complex traits

We demonstrate clinical relevance of the improved genotyping and imputation through an application of coronary artery disease (CAD) polygenic risk scores (PRS) in an independent South Asian cohort (1800 cases, 1163 controls) which were genotyped on the GSA3 arrays^22^. We imputed the genotypes using the 1000 Genomes and GAsP2 panels and applied the ancestry-adjusted genome-wide PRS model from ref. 22. The results showed a marked improvement in the predictive power of the PRS, with an improved AUC (0.638 for GASP2 vs. 0.595 for 1000 Genomes). The odds ratios of CAD for individuals in the top deciles (9th-10th) compared to those in the middle deciles (5th-6th) are higher in the GAsP2-based PRSs (OR_9th_ = 1.67; OR_10th_ = 2.43) as compared to the 1000 Genomes-based PRSs (OR_9th_ = 1.32; OR_10th_ = 1.83; Fig 4c). These improved ORs are on par with those achieved for European samples with the appropriate imputation panel (HRC, UK10K, 1000 Genomes), GWAS, and PRS model (UKB OR_9th_ = 1.51; OR_10th_ = 2.52; Fig 4c). This improved performance can be explained by the improved imputation accuracy as well as the increased number of well-imputed variants (Supplementary Table 3).

## Conclusion

South Asian populations provide a rich potential for human genetic discovery that is largely unexplored. A population-scale genotyping project in South Asia will open up opportunities to explore disease genetics in ways that are impractical or infeasible in other populations. Notably, the dramatically higher rate of homozygosity that is found in parts of South Asia allows homozygous loss of function effects to be studied for many genes that cannot realistically be accessed in outbred populations such as those that predominate in Europe and East Asia^17,23^. Traditionally, homozygous gene function has been explored through family-based studies, often involving self-identified consanguineous unions. Although this will continue to be an effective way to carry out focused studies, a population-scale dataset in South Asia will facilitate identification of appropriate families and will also open up new opportunities to consider homozygosity in population-based association analyses. At the same time, this dataset will provide the opportunity to evaluate disease associations with a novel set of functional variants, *e*.*g*. the unique set and larger set of pLoF alleles with frequencies > 0.1% found in South Asians as compared to Europeans. The SARGAM genotyping array and the GAsP2 imputation reference panel allow South Asian genotypes to be captured in an economical and effective manner.

Population-based genetic studies have been effectively carried out within single coordinated health-care delivery systems such as national single-payer systems. South Asia provides a different set of challenges and a different set of possibilities. In India in particular, Super-specialty hospitals, organized to deliver health-care in a specific disease area in a way that takes advantage of the economies of scale presented by its large population base, predominate in certain markets. These hospitals can, in a disease focused fashion, rival the scale of national general hospital systems of some countries. Thus while, national biobank systems do not exist in South Asia at present that could provide a foundation for a broad cross-sectional evaluation of the genetics of disease, the scale at which patients can be recruited within specific disease areas will allow clinically relevant datasets to be constructed to a total size that is unrealistic in most parts of the world. This, paired with the unique population structure of South Asia, presents a powerful set of opportunities for genetic discovery that will improve healthy life-span around the globe.

## Methods

### Samples

We utilized a combination of genomes from previously published studies^7-12^, newly sequenced genomes from 1000 Genomes Project samples, and newly sequenced genomes from several ongoing genetic studies in South Asia. Further information on the samples is contained in Supplementary Information 1 and Supplementary Table 1.

### Sequencing, filtering, alignment and variant calling

Illumina short reads were mapped to the reference genome (build GRCh38) using BWA^24^. We then used GATK4^25^ for base quality score recalibration, indel realignment, duplicate removal, variant discovery and joint genotyping, using the GATK Best Practices recommendations^26,27^. We then removed variants that were monomorphic or were not annotated as PASS, and converted genotype calls with genotype quality (GQ) score < 20 to missing data. Finally, we removed any variants with a missing genotype rate of > 30%.

### Comparison with 1000 Genomes Data

We downloaded genotype calls from the high-coverage 1000 Genomes Project data from http://ftp.1000genomes.ebi.ac.uk/vol1/ftp/data_collections/1000G_2504_high_coverage/working/20190425_NYGC_GATK/. We then used the same filters as described above, except with a genotype quality filter of > 40. Then, for 22 individuals that were sequenced independently but were contained in both our call set and the 1000 Genomes Project (high-coverage) call set, we tabulated the non-reference discordance rate of the filtered genotype calls in the two data sets. Results are summarized in Supplementary Table 2.

### Sample QC and identification of 1^st^ degree relative pairs

We used KING^28^ to identify close relatives in our data. We labeled pairs of individuals as duplicates or 1st degree relatives if the estimated kinship coefficients were > 0.4 and [0.177, 0.4] respectively. We then removed samples in the following order:

1. All samples with duplicates from another population
2. For remaining duplicate pairs in the same population, the individual with more missing data
3. Individuals that have more than one 1st degree relative
4. Individuals with genotype calls at < 90% of all SNPs
5. For remaining 1st degree relative pairs, the individual with more missing data

After this filtering, we were left with 6,443 genomes for downstream analyses.

### Phasing

The collection of 6,443 individuals described above was computationally phased using eagle2^29^, with the default options (which includes allowing eagle2 to impute sporadic missing genotypes). The GRCh38-based genetic map made available with the eagle2 distribution was used for the phasing. We also used the same workflow to construct a reference panel consisting of only the South Asian medical cohorts, which was used in the design of the SARGAM array.

### Population structure (Fst, PCA, UMAP, and Admixture)

We used *plink* version 1.9^30^ to conduct Principal Components analysis. We filtered SNPs to have a MAF > 0.01, and LD-pruned using an R^2^ threshold of 0.2. We then created PCA plots after removing the pruned SNPs with the *variant-weights* modifier in *plink1*.*9*. These analyses were performed separately for different groups of individuals after the removal of 1st degree relative pairs and low-coverage samples as described above.

UMAP projection was performed using the protocol and script published by Diaz-Papkovich and colleagues^31^. 15 principal components were used to generate the two dimensional UMAP projection. Based on visualization and separation of known population groups in the Birbhum Cohort, we chose the key parameter settings as follows: number of neighbors (NN) of 15 and minimum distance (MD) of 0.5.

The ADMIXTURE^14^ analysis was performed using Version 1.3.0 of the software (http://www.genetics.ucla.edu/software/admixture). We used SNPs with MAF > 0.01, with the call rate > 99.9%, and LD-pruned using a 50-SNP sliding window and variance inflation factor threshold of 2. The number of components K was optimized to minimize the cross-validation error using Chromosome 21. The optimal K for the Birbhum Cohort was 4, but for the larger South Asian and global samples, the cross-validation error continued to decrease even for a large K (K=40). Thus, we chose to present the results at K=12 which is the same number of components used in the GenomeAsia 100K Pilot study^12^.

Weir and Cockerham weighted F_ST_ estimates were calculated using VCFtools^32^ Version 0.1.17. Only MAF-filtered and LD-pruned markers, as described above, were used. Samples that were related or did not pass QC were excluded.

### Rare homozygotes

For each population considered, we stratified variants according to the MAF in the specific population and tabulated the number of rare homozygotes and the expected number of rare homozygotes for each MAF category, assuming random mating. We then further stratified these results (Fig. 2c-d) by classifying some individuals as “inbred” (i.e., offspring of 3^rd^ degree relatives or closer) or “outbred” (i.e., offspring of 6^th^ degree relatives or more distant), using the estimation process described below.

### Runs of Homozygosity (ROH)

We used PLINK version 1.9^30^ to identify ROH in our data. We used the default parameter settings, except for the following:

--homozyg --homozyg-kb 500 --homozyg-window-snp 100 --homozyg-window-het 2 --homozyg-window-missing 20 --maf 0.001

We then converted the lengths of all ROHs into genetic distances, using a genetic map first created by Adam Auton and downloaded from the Beagle website at https://bochet.gcc.biostat.washington.edu/beagle/genetic/maps/

We required ROHs to have a minimum physical length of 500 Kb and a minimum genetic distance of 1 cM to be included in our analyses.

### Estimating the degree of inbreeding

We used a summary likelihood approach for estimating the degree of inbreeding from an individual’s ROH tracts. We focused on the longest ROHs to provide better power for distinguishing ROHs that arise due to endogamy versus ones that arise due to very recent inbreeding. Specifically, we tabulated (1) the number of ROHs longer than 10 cM, and (2) the sum of the genetic lengths of the 10 longest ROHs for each individual. Simulations suggest that these two summaries are more informative than other, similar summaries based on the number or length of the longest ROH tracts (Results not shown.) Concurrently, we simulated the distribution of ROH lengths expected under various degrees of inbreeding (see section directly below), ranging from the offspring of 2^nd^ degree relatives (e.g., uncle - niece) to the offspring of 6^th^ degree relatives (e.g., half 2^nd^ cousins). Then, for each individual, we estimated the probability of observing the two ROH summaries (within 1% for the 10 longest ROHs) as a function of the degree of inbreeding. We then assigned the degree of inbreeding with the highest likelihood for each individual, treating 6^th^ degree relatives to be “unrelated”.

### Simulating expected ROH size distribution under inbreeding

We utilize the model of Clark^33^ for simulating the distribution of autozygous segments expected under a specific degree of inbreeding. Specifically, we assume that the genetic lengths of chromosomal segments inherited from particular paternal and maternal ancestors follows an exponential distribution with mean equal to 100 cM divided by the total number of generations in the path from the proband back to the particular ancestors. For example, for an individual whose parents are 1st cousins, there are four paternal great grandparents (and thus eight total paternal autosomal chromosomes three generations ago) and eight total maternal autosomal chromosomes that could be inherited at any particular genomic location. Of the 8 x 8 = 64 possible inheritance patterns, four result in consanguinity. We model an autosome’s ancestry as a series of blocks of ancestry, each with genetic length exponentially distributed with mean 100 / 6 cM, and with each block having a 4 / 64 = 6.25% chance of being autozygous. We take the genetic lengths of chromosomes from the original deCODE genetic map^34^, and tabulate the number and size distribution of autozygous segments over 2 million simulations for each degree of consanguinity considered.

### Loss of Function variants

A list of high-confidence Loss-of-Function (LoF) variants, including frameshift, splice-site, non-sense, start-loss and stop-loss mutations, was obtained using the following criteria:

- The LoF variants should be predicted as high-confidence (HC) from the LOFTEE program^35^
- The LoF variants must fall within the high-confidence regions defined by the Genome-In-a-Bottle^36^ (GIAB) consortium (version - v.3.3.2)
- The LoF variants cannot fall in segmental duplication regions of the genome (genomicSuperDups) as defined by the UCSC Genome Browser
- The Ensembl/GENCODE transcript with highest expression among all the transcripts of the gene was retained. The expression value was obtained from the GTEx (Genotype-Tissue Expression) project (version 8) and averaged over all median tissue expressions.

### Burden test and association test

We analyzed a total of 2,994 South Indian samples for which we had exome or whole-genome sequence data as well as blood lipid level measurements. Samples with extreme blood lipid values, defined as values outside of Q1 - 1.5 IQR/sqrt(n) and Q3 + 1.5 IQR/sqrt(n), were removed. Variant annotation was carried out using the Variant Effect Predictor^37^ annotated against Ensembl v75^38^. We considered only the LoF variants using the filters described above. Further, we removed samples and variants with poor call rates (>5% no calls) and only kept the variants with MAF < 0.1. For each sample, we combined the LoF variant dosages into a single burden within each gene and restricted the analysis to genes with at least 3 LoF variant carriers. Association analyses of quantitative traits were perfromed using linear regression on the burden score with age and sex as covariates.

### SARGAM array design

We partnered with Thermo Fisher Scientific to develop a custom genotyping array using their Axiom platform. The SARGAM (South Asian Research Genotyping Array for Medicine) assays a total of 639,029 SNPs, including 515,921 variants chosen to optimize imputation accuracy as well as 102,752 putatively functional variants that had a minor allele frequency of > 0.1% in our medical cohorts. The imputation-based SNPs were chosen using an algorithm similar to the one described by Hoffmann and colleagues^39^, based on the South Asian medical cohort phased reference panel described above. The initial list of putative functional variants were obtained from a variety of sources, including this project, gnomAD^17^, the UK Biobank^40^ and properly consented MedGenome internal data.

We initially started with a larger list of 924,667 SNPs that were assayed on two custom test arrays that were then used to genotype 960 individuals from the GAsP2 study for quality control purposes. We removed SNPs that could not be genotyped accurately as well as low-priority variants to arrive at the final SARGAM array design.

### Polygenic Risk Scores

#### South Asian samples

We used the genotype data of 1,800 CAD cases and 1,163 controls assayed using Illumina GSA3 array covering more than 600,000 genome-wide markers, as reported in ref. 22. All the samples had more than 95% of the markers successfully genotyped. We used Beagle5.0^41^ to impute all variants with minor allele count > 4, using either the 1000 Genomes Project Phase 3 data or the GAsP2 reference panel described above. The total number of imputed variants were 24,154,211 and 24,969,892 respectively. We used the markers reported by the CardiogramplusC4D consortium^42^, and the methods described in ref. 22 to construct polygenic risk scores.

#### UK Biobank European samples

We selected 2910 samples from the UK Biobank, comprising 1448 CAD cases and 1468 controls having European ancestry. The cases were selected with ICD-9 codes of 410.X, 411.0, 412.X, 429.79 or ICD-10 codes of I21.X, I22.X, I23.X, I24.1, I25.2. We used the imputed genetic data for generation of polygenic risk scores. This research was conducted using the UK Biobank Resource under Application Number 42406.

## References

1. Norio, R. Finnish Disease Heritage I: characteristics, causes, background. Hum. Genet. 112, 441–456 (2003).

2. Gross, S. J., Pletcher, B. A., Monaghan, K. G., & Professional Practice and Guidelines Committee. Carrier screening in individuals of Ashkenazi Je ish descent. Genet. Med. 10, 54–56 (2008).

3. Payne, M., Rupar, C. A., Siu, G. M., & Siu, V. M. Amish, mennonite, and hutterite genetic disorder database. Paediatr. Child Health 16, e23–e24 (2011).

4. Gudb artsson, D. F. et al. Large-scale hole-genome sequencing of the Icelandic population. Nat. Genet. 47, 435–444 (2015).

5. Reich, D. et al. Reconstructing Indian population history. Nature 461, 489–494 (2009).

6. Mastana, S. S. Unity in diversity: an overvie of the genomic anthropology of India. Ann. Hum. Biol. 41, 287–299 (2014).

7. Wong, L.-P. et al. Deep hole-genome sequencing of 100 southeast Asian Malays. Am. J. Hum. Genet. 92, 52–66 (2013).

8. Wong, L.-P. et al. Insights into the genetic structure and diversity of 38 South Asian Indians from deep hole-genome sequencing. PLoS Genet. 10, e1004377 (2014).

9. Vernot, B. et al. Excavating Neandertal and Denisovan DNA from the genomes of Melanesian individuals. Science 352, 235–239 (2016).

10. Lu, D. et al. Ancestral origins and genetic history of Tibetan highlanders. Am. J. Hum. Genet. 99, 580–594 (2016).

11. Mallick, S. et al. The Simons Genome Diversity Pro ect: 300 genomes from 142 diverse populations. Nature 538, 201–206 (2016).

12. GenomeAsia 100K Consortium. The GenomeAsia 100K pro ect enables genetic discoveries across Asia. Nature 576, 106–111 (2019).

13. Price, A. L., et al. Principal components analysis corrects for stratification in genomeide association studies. Nat. Genet. 38, 904–909 (2006).

14. Alexander, D. H., Novembre, J., & Lange, K. Fast model-based estimation of ancestry in unrelated individuals. Genome Res. 19, 1655–1664 (2009).

15. McInnes, L., Healy, J., & Melville, J. UMAP: Uniform Manifold Approximation and Pro ection for dimension reduction. arXiv, available at https://arxiv.org/abs/1802.03426 (2018).

16. Nakatsuka, N. et al. The promise of discovering population-specific disease-associated genes in South Asia. Nat. Genet. 49, 1403–1407 (2017).

17. Karcze ski, K. J. et al. The mutational constraint spectrum quantified from variation in 141,456 humans. Nature 581, 434–443 (2020).

18. Tall, A. R. Functions of cholesterol ester transfer protein and relationship to coronary artery disease risk. J. Clin. Lipidol. 4, 389–393.

19. Tarugi, P., Bertolini, S., & Calandra, S. Angiopoietin-like protein 3 (ANGPTL3) deficiency and familial combined hypolipidemia. J. Biomed. Res. 33, 73–81.

20. The TG and HDL orking group of the Exome Sequencing Pro ect, NHLBI. Loss-of-function mutations in APOC3, triglycerides, and coronary disease. N. Engl. J. Med. 371, 22–31.

21. Saleheen, D. et al. Human knockouts and phenotypic analysis in a cohort ith a high rate of consanguinity. Nature 544, 235–239 (2017).

22. Wang, M. et al. Validation of a genomeide polygenic score for coronary artery disease in South Asians. J. Am. Coll. Cardiol. 76, 703–714 (2020).

23. Lek, M. et al. Analysis of protein-coding genetic variation in 60,706 humans. Nature 536, 285–291 (2016).

24. Li, H., & Durbin, R. Fast and accurate short read alignment ith Burro s-Wheeler transform. Bioinformatics 25, 1754–1760 (2009).

25. McKenna, A. et al. The Genome Analysis ToolKit: a MapReduce frame ork for analyzing next-generation DNA sequencing data. Genome Res. 20, 1297–1303 (2010).

26. DePristo, M. A. et al. A frame ork for variation discovery and genotyping using next-generation DNA sequencing data. Nat. Genet. 43, 491–498 (2011).

27. Van der Au era, G. A. et al. From FastQ data to high-confidence variant calls: the Genome Analysis ToolKit best practices pipeline. Curr. Protoc. Bioinformatics 43, 11.10.1-11.10.33 (2013).

28. Manichaikul, A. et al. Robust relationship inference in genomeide association studies. Bioinformatics 26, 2867–2873 (2010).

29. Loh, P.-R. et al. Reference-based phasing using the Haplotype Reference Consortium panel. Nat. Genet. 48, 1443–1448 (2016).

30. Purcell, S. et al. PLINK: a tool set for hole-genome association and population-based linkage analyses. Am. J. Hum. Genet. 81, 559–575 (2007).

31. Diaz-Papkovich, A., Anderson-Trocme, L., Ben-Eghan, C., & Gravel, S. UMAP reveals cryptic population structure and phenotype heterogeneity in large genomic cohorts. PLoS Genet. 15, e1008432 (2019).

32. Danacek, P. et al. The variant call format and VCFtools. Bioinformatics 27, 2156–2158 (2011).

33. Clark, A. G. The size distribution of homozygous segments in the human genome. Am. J. Hum. Genet. 65, 1489–1492 (1999).

34. Kong, A. et al. A highresolution recombination map of the human genome. Nat. Genet. 31, 241–247 (2002).

35. LOFTEE (Loss Of Function Transcript Effect Estimator). Available from https://github.com/konradk/loftee

36. Zook, J. M. et al. Extensive sequencing of seven human genomes to characterize benchmark reference materials. Scientific Data 3, 160025 (2016).

37. McLaren, W. et al. The Ensembl Variant Effect Predictor. Genome Biol. 17, 122 (2016).

38. Yates, A. D. et al. Ensembl 2020. Nucleic Acids Res. 48, D682–D688 (2020).

39. Hoffmann, T. J. et al. Next generation genomeide association tool: design and coverage of a high-throughput European-optimized SNP array. Genomics 98, 79–89 (2011).

40. Bycroft, C. et al. The UK Biobank resource ith deep phenotyping and genomic data. Nature 562, 203–209 (2018).

41. Bro ning, B. L., Zhou, Y., & Bro ning, S. R. A one-penny imputed genome from next generation reference panels. Am. J. Hum. Genet. 103, 338–348 (2018).

42. Nikpey, M. et al. A comprehensive 1000-Genomes based genomeide association meta-analysis of coronary artery disease. Nat. Genet. 47, 1121–1130 (2015).

